# Homology-guided re-annotation improves the gene models of the alloploid *Nicotiana benthamiana*

**DOI:** 10.1101/373506

**Authors:** Jiorgos Kourelis, Farnusch Kaschani, Friederike M. Grosse-Holz, Felix Homma, Markus Kaiser, Renier A. L. van der Hoorn

## Abstract

*Nicotiana benthamiana* is an important model organism of the Solanaceae (Nightshade) family. Several draft assemblies of the *N. benthamiana* genome have been generated, but many of the gene-models in these draft assemblies appear incorrect. Here we present an improved re-annotation of the Niben1.0.1 draft genome assembly guided by gene models from other *Nicotiana* species. This approach overcomes problems caused by mis-annotated exon-intron boundaries and mis-assigned short read transcripts to homeologs in polyploid genomes. With an estimated 98.1% completeness; only 53,411 protein-encoding genes; and improved protein lengths and functional annotations, this new predicted proteome is better than the preceding proteome annotations. This dataset is more sensitive and accurate in proteomics applications, clarifying the detection by activity-based proteomics of proteins that were previously mis-annotated to be inactive. Phylogenetic analysis of the subtilase family of hydrolases reveal a pseudogenisation of likely homeologs, associated with a contraction of the functional genome in this alloploid plant species. We use this gene annotation to assign extracellular proteins in comparison to a total leaf proteome, to display the enrichment of hydrolases in the apoplast.

## INTRODUCTION

*Nicotiana benthamiana* has risen to prominence as a model organism for several reasons. First, *N. benthamiana* is highly susceptible to viruses, resulting in highly efficient virus-induced gene-silencing (VIGS) for rapid reverse genetic screens (Senthil-Kumar and Mysore, 2014). This hypersusceptibility to viruses is due to an ancient disruptive mutation in the RNA-dependent RNA polymerase 1 gene (*Rdr1*), present in the lineage of *N. benthamiana* which is used in laboratories around the world (Bally *et al.*, 2015). Reverse genetics using *N. benthamiana* have confirmed many genes important for disease resistance (Wu *et al.*, 2017; Senthil-Kumar *et al.*, 2018). Second, *N. benthamiana* is highly amenable to the generation of stable transgenic lines (Clemente, 2006; Sparkes *et al.*, 2006) and to transient expression of transgenes (Goodin *et al.*, 2008). This easy manipulation has facilitated rapid forward genetic screens and has established *N. benthamiana* as the plant bioreactor of choice for the production of biopharmaceuticals (Stoger *et al.*, 2014). Finally, *N. benthamiana* is a member of the Solanaceae (Nightshade) family which includes important crops such as potato (*Solanum tuberosum*), tomato (*Solanum lycopersicum*), eggplant (*Solanum melongena*), and pepper (*Capsicum* ssp.), as well as tobacco (*Nicotiana tabacum*) and petunia (*Petunia* ssp.).

*N. benthamiana* belongs to the *Suaveolentes* section of the *Nicotiana* genus, and has an ancient allopolyploid origin (∼6Mya, Clarkson et al., 2017) accompanied by chromosomal re-arrangements resulting in a complex genome with 19 chromosomes per haploid genome, a reduced number when compared to the ancestral allotetraploid 24 chromosomes. The estimated haploid genome size is ∼3.1Gb (Goodin et al., 2008; Leitch et al., 2008; Wang and Bennetzen, 2015). There are four independent draft assemblies of the *N. benthamiana* genome (Bombarely *et al.*, 2012; Naim *et al.*, 2012), as well as a *de*-*novo* transcriptome generated from short-read RNAseq (Nakasugi *et al.*, 2014). These datasets have greatly facilitated research in *N. benthamiana*, allowing for efficient prediction of off-targets of VIGS (Fernandez-Pozo *et al.*, 2015) and genome editing using CRISPR/Cas9 (Liu *et al.*, 2017), as well as RNAseq and proteomics studies (Grosse-Holz *et al.*, 2018).

During our research using proteomics and reverse genetic approaches, we realized that many of the gene models in these draft assemblies are incorrect, and that putative pseudo-genes are often annotated as protein-encoding genes. This is at least partly because these draft assemblies are highly fragmented and because *N. benthamiana* is an old alloploid. Mapping short read sequences onto polyploid draft genomes frequently results in mis-assigned transcripts and mis-annotated gene models (Vaattovaara et al., 2019). The *de-novo* transcriptome assembly also has a high proportion of chimeric transcripts. Because of incorrect annotations, extensive bioinformatics analysis is required to select target genes for reverse genetic approaches such as gene silencing and editing, or for phylogenetic analysis of gene families.

We realized that the genome annotations of several other species in the *Nicotiana* genus generated using the NCBI Eukaryotic Genome Annotation Pipeline were much better, and we therefore decided to re-annotate the available *N. benthamiana* draft genomes using these *Nicotiana* gene models as a template. We generated a full genome annotation of the protein-encoding genes in the Niben1.0.1 draft genome assembly and extracted gene-models missing from this annotation from the other draft genome assemblies. Here we show that this dataset explains activity-based proteomics datasets better, is more accurate and sensitive for proteomics, and that this annotation greatly facilitates phylogenetic analysis of gene families.

## RESULTS AND DISCUSSION

### Re-annotation of gene-models in the *N. benthamiana* genome assemblies

The observation that predicted proteins from other *Nicotiana* species were more correct for several gene families inspired us to use the corresponding coding sequences to re-annotate the *N. benthamiana* genome. We used Scipio (Keller et al., 2008) to transfer the gene models from *Nicotiana* species to *N. benthamiana*. Scipio refines the transcription start-site, exon-exon boundaries, and the stop-codon positions of protein-encoding sequences aligned to the genome using BLAT (Keller *et al.*, 2008). When input protein-encoding sequences are well-annotated, this method is more accurate and sensitive than other gene prediction methods (Keller *et al.*, 2011).

To generate the input dataset, we selected the predicted protein sequences from recently sequenced *Nicotiana* species (Sierro et al., 2013, 2014; Xu et al., 2017) generated using the NCBI Eukaryotic Genome Annotation Pipeline (**Figure 1**) and used CD-HIT at a 95% identity cut-off to reduce the redundancy and remove partial sequences (**Figure 1**, Step 1). The resulting dataset, Nicotiana_db95, contains 85,453 protein-encoding sequences from various *Nicotiana* species. We then used this Nicotiana_db95 dataset as an input dataset to annotate gene-models in the Niben1.0.1 draft genome assembly using Scipio (**Figure 1**, Step 2).

**Figure 1.**
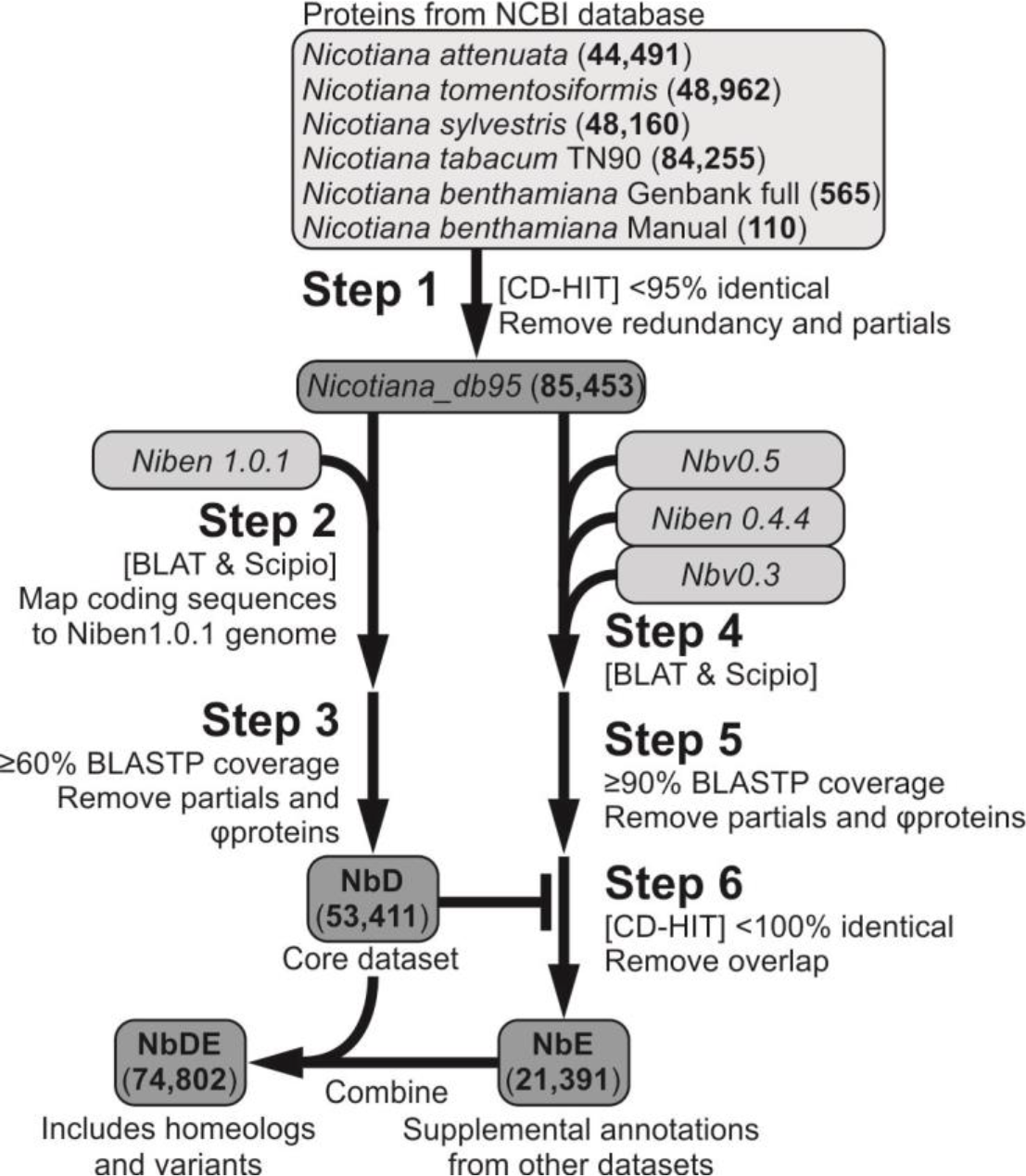
Bioinformatics pipeline for improved *Nicotiana benthamiana* proteome annotation. The predicted proteins of *Nicotiana* species generated by the NCBI Eukaryotic Genome Annotation Pipeline were retrieved from Genbank and clustered at 95% identity threshold to reduce redundancy (**Step 1**), and used to annotate the Niben1.0.1 genome assembly (**Step 2**). Only those proteins with an alignment coverage ≥60% to the *Nicotiana* predicted proteins as determined by BLASTP were retained (**Step 3**) to produce the NbD core dataset. Similarly, the other draft genome assemblies were annotated (**Step 4**), and only those proteins with an alignment coverage ≥90% to the *Nicotiana* predicted proteins as determined by BLASTP were retained (**Step 5**). CD-HIT-2D was used at 100% sequence identity to retain proteins missing in NbD dataset (**Step 6**), resulting in supplemental dataset NbE. NbD and NbE can be combined (NbDE) to maximise the spectra annotation for proteomics experiments.

The Scipio-generated dataset contains many full-length protein-encoding sequences, but also partial sequences caused by fragmentation in the Niben1.0.1 dataset. In addition, we noticed many incomplete gene products caused by premature stop-codons and frameshift mutations. To remove these partial sequences and putative pseudogene products from the protein dataset, we used BLASTP against the NCBI *Nicotiana* reference proteomes and retained only protein-encoding sequences with ≥60% coverage. This resulted in 53,411 protein-encoding gene annotations (NbD dataset, **Figure 1**, Step 3).

To complement the NbD dataset, we used the same approach to re-annotate the Nbv0.5, Niben0.4.4 and Nbv0.3 draft assemblies using Scipio and the Nicotiana_db95 dataset as template (Step 4). We retained proteins with ≥90% BLASTP coverage with the *Nicotiana* reference proteomes in NCBI (**Figure 1**, Step 5) and we used CD-HIT-2D at a 100% threshold to select coding sequences not present in the NbD dataset (Step 6). The resulting supplemental NbE dataset contains 21,391 additional protein-encoding sequences (**Figure 1**). Besides many new gene models, this NbE dataset will also contain sequences that are nearly identical to those in the NbD database. These proteins may be derived from homeologs caused by polyploidisation, but can also be caused by sequence polymorphisms between different sequenced plants and by sequencing and assembly errors. For protein annotation in MS experiments, however, the combination of the NbD and NbE datasets (the NbDE dataset) will maximise the annotation of the spectra.

### The NbDE proteome is more complete, sensitive, accurate, and relatively small

We next compared the predicted proteome to the published predicted proteomes. The NbD and NbDE datasets have relatively few entries when compared to the other datasets (**Figure 2A**). The preceding datasets include the predicted proteomes from the Niben0.4.4 (76,379 entries, Bombarely *et al.*, 2012)) and Niben1.0.1 (57,140 entries, Bombarely *et al.*, 2012)). We also included a previously described curated database in which gene-models from Niben1.0.1 were corrected using RNAseq reads (74,093 entries, Grosse-Holz *et al.*, 2018)), and the predicted proteome (Nbv5.1*) derived from a *de-novo* transcriptome (191,039 entries, Nakasugi *et al.*, 2014). We also included the Nicotiana_db95 dataset (85,453 entries) in this comparison.

**Figure 2.**
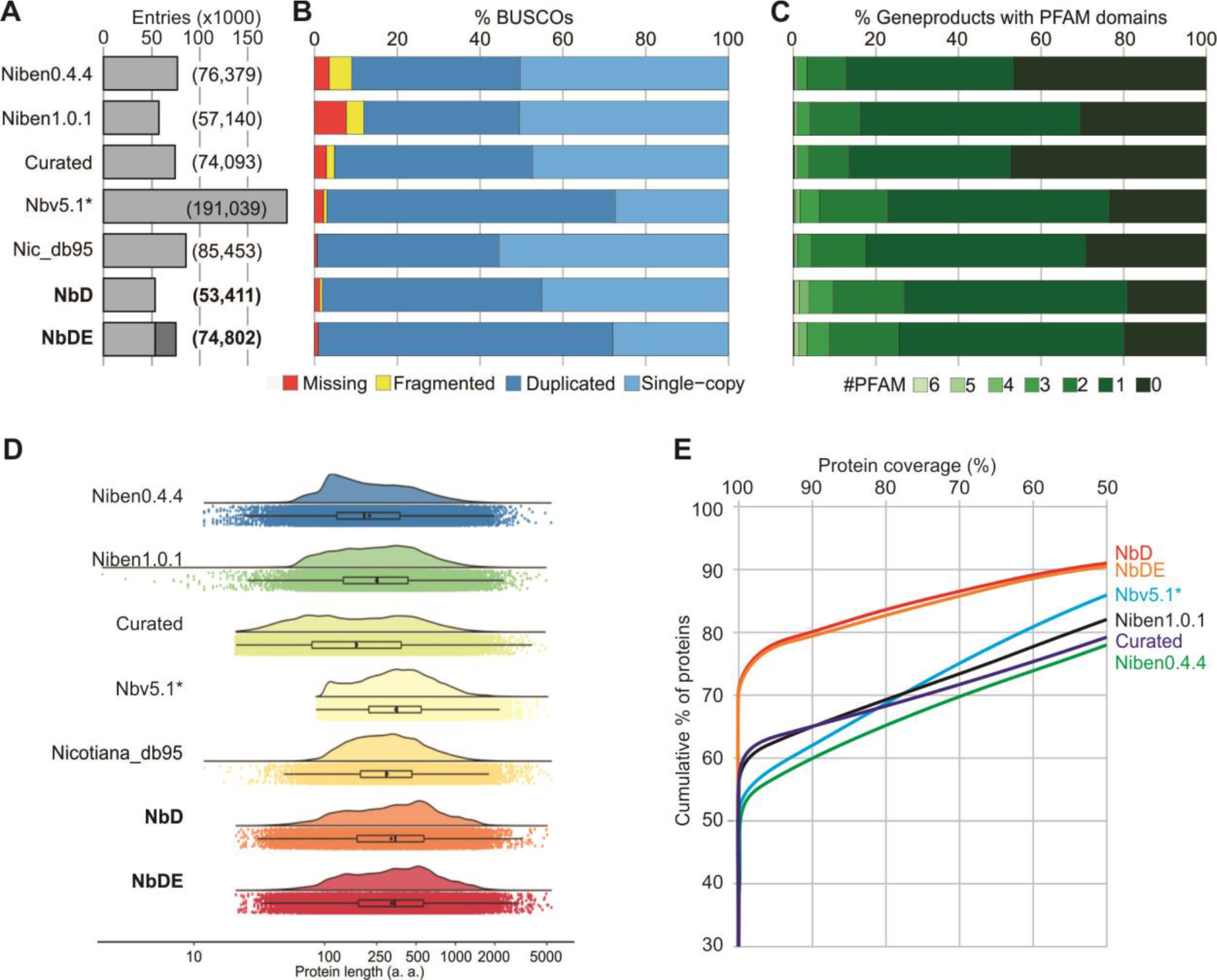
Increased lengths, coverage and annotation of *N. benthamiana* proteins. **(A)** NbD/NbDE datasets have relatively few entries when compared to preceding datasets. **(B)** NbD/NbDE datasets contain nearly all benchmark genes as full-length genes, according to Benchmarking Universal Single-Copy Orthologs (BUSCO) of embryophyta. **(C)** The NbD/NbDE datasets have higher number of annotated PFAM domains. **(D)** NbD/NbDE datasets have relatively longer protein lengths. Violin and boxplot graph of log_10_ protein length distribution of each database. Jittered dots show the raw underlying data. **(E)** NbD/NbDE annotated proteins have a higher percentage coverage to the tomato proteins as determined by BLASTP.

To determine the completeness of the NbD and NbDE datasets, we analysed the presence of BUSCOs (Benchmarking Universal Single-Copy Orthologs (Simão *et al.*, 2015; Waterhouse *et al.*, 2018)). We used the embryophyta BUSCO set, which contains 1440 highly conserved plant genes that are predominantly found as single-copy genes (Simão *et al.*, 2015). Nicotiana_db95 has one fragmented and nine missing BUSCOs, indicating that this dataset contains 99.3% of the *Nicotiana* genes (**Figure 2A**). In the NbD dataset, 98.1% of the BUSCO proteins were identified as full-length proteins, and only 0.6% of the BUSCOs is fragmented and 1.3% is missing (**Figure 2A**). The 98.1% full length BUSCO proteins in the NbD dataset consists of 45% single-copy genes and 53.1% duplicated genes, consistent with the allotetraploid nature of *N. benthamiana*. The combined NbDE dataset contains 99.0% of the BUSCOs and contains 71.1% duplicated full length BUSCOs (**Figure 2A**), but part of this increased duplication may be due to small sequence variations between the different genome assemblies rather than from genes missing in the NbD annotation. The BUSCO scores of the NbD and NbDE datasets are superior when compared to previously published annotations. The best previously predicted proteome is from dataset Nbv5.1, which has 96.9% of the BUSCOs as full-length proteins, whereas 0.8% is missing and 2.2% is incomplete (**Figure 2A**). However, Nbv5.1 contains nearly five times more protein coding sequences than NbD and 69.7% of BUSCOs are duplicated.

Second, we investigated the number of unique PFAM identifiers found with each entry in each proteome, because mis-annotated sequences and fragmented gene products are less likely to receive a PFAM annotation (Vaattovaara et al., 2019). Indeed, over 80% of the proteins in the NbD and NbDE datasets get at least one PFAM identifier whereas this match is lower with the preceding datasets, indicating that proteins in NbD and NbDE are better annotated (**Figure 2B**).

Third, we mapped the length distributions of the predicted proteins in comparison with the Nicotiana_db95 dataset. The proteins in the NbD and NbDE datasets are significantly longer than those in the previously predicted proteomes except for the Nbv5.1 primary + alternate proteome (**Figure 2C**). This is probably because the Niben0.4.4, Niben1.0.1 and curated datasets contain many partial genes and pseudo-genes, while the Nbv5.1 primary + alternative proteome has a high proportion of chimeric sequences, which are biased towards long transcripts due to the assembly of short sequencing reads.

Fourth, to verify that the increased protein lengths is not due to incorrect annotation, we performed BLASTP searches against the *Solanum lycopersicum* RefSeq proteins (annotation v103), using the *N. benthamiana* annotated proteins as the query and the *S. lycopersicum* RefSeq proteins as the target. 70% of the NbD and NbDE proteins show a full coverage of tomato proteins, whereas in the other datasets less than 60% of the proteins show full coverage (**Figure 2D**). This indicates that in addition to a more complete annotation with fewer missing genes as determined by BUSCO, the NbD and NbDE datasets are also more accurate in the annotation of gene models.

Finally, since phylogenetic analysis of gene families in closely related species often relies on gene-annotations, we compared our NbD proteome annotation against the predicted proteomes of Solanaceae species for which genomes have been sequenced (Supplemental **Figure S1**). Our NbD proteome compare well to the predicted proteomes of other sequences Solanaceae species generated by the NCBI Eukaryotic Genome Annotation Pipeline. In addition, since the predicted proteomes of some of these species were not annotated using the NCBI pipeline, they miss a relatively high proportion of genes (up to 28.5% of genes missing or fragmented). Therefore, care must be taken to not over-interpret results derived from phylogenetic analysis using these datasets.

### Improved annotation of spectra in proteomics experiments

We next tested the annotation of MS spectra from four biological replicates of total leaf extracts using the different datasets. Both the NbD and NbDE datasets outperform the preceding datasets (Figure 3A). This is notable because the NbD dataset has the fewest entries and yet works well in spectra annotation. More specifically, the NbD dataset also identifies the highest number of unique peptides per protein, consistent with having the fewest entries and increased length (Figure 3B). These metrics show that the new NbD and NbDE datasets are more sensitive and accurate for proteomics than the currently available datasets.

**Figure 3.**
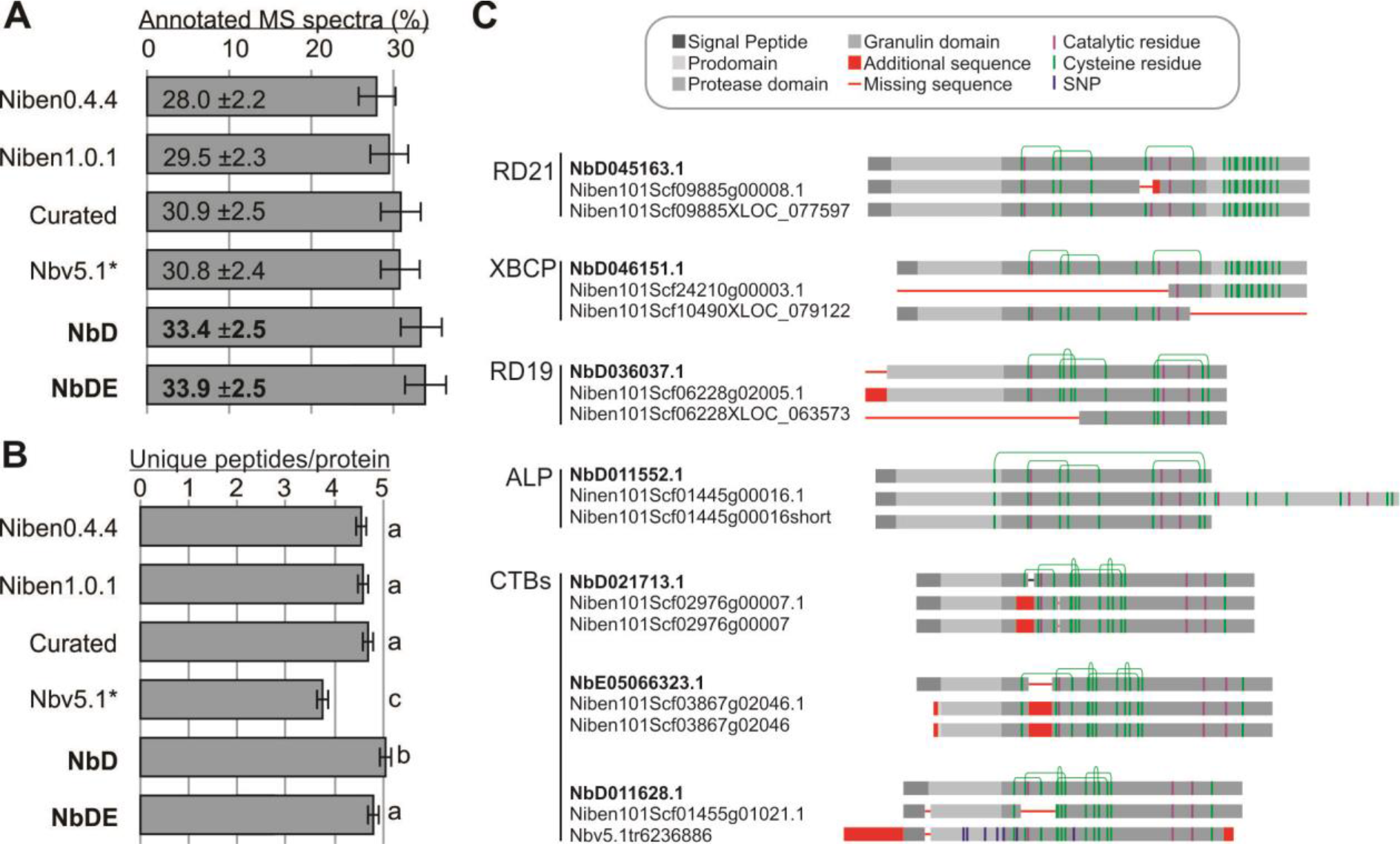
NbD/NbDE datasets outperform the annotation of spectra in proteomics. **(A)** Percentage of annotated MS/MS spectra in total leaf extract samples. **(B)** Average number of unique peptides assigned per protein in the different databases. **(A-B)** Means and standard error of the mean are shown for four biological replicates of total leaf extracts. **(C)** Mis-annotations of papain-like Cys proteases (PLCPs) detected by activity-based protein profiling (Jutras et al., 2019). Leaf extracts were labelled with activity-based probes for PLCPs and labelled proteins were purified and analysed by MS. Shown are the protein annotations found in the NbDE (top) Niben1.0.1 (middle) and curated datasets (bottom), highlighting mis-annotations (red) caused by partial transcripts, mis-annotation of exon-intron boundaries, and mis-assemblies.

We independently validated the NbD and NbDE annotations on an independent dataset where we re-analysed a previously published apoplastic proteome of agro-infiltrated *N. benthamiana* as compared to non-infiltrated *N. benthamiana* (PRIDE repository PXD006708, Grosse-Holz et al., 2018). Both the NbD and NbDE annotations are more sensitive and accurate than the curated dataset on this experiment (18,059 and 18,352 vs 17,954 peptides detected; 21.85% ±3.0% and 22.2% ±3.1% vs 21.7% ±3.0% spectra identified). This independently confirms the high performance of the NbD and NbDE datasets.

To confirm that our new annotations are correct and biologically relevant we examined the annotation of 14 papain-like Cys proteases (PLCPs) that we identified recently from agroinfiltrated *N. benthamiana* leaves (Jutras *et al.*, 2019). These proteases were active proteins because they reacted with an activity-based probe (DCG-04, a biotinylated PLCP inhibitor) to facilitate the purification and detection of these proteins. Of the 14 detected PLCPs, eight were identical between the datasets. However, six PLCPs were mis-annotated in the Niben1.0.1 and curated datasets (Figure 3C). Two mis-annotated PLCPs lacked catalytic residues; one PLCP lacked a Cys residue that is crucial for PLCP stability; two PLCPs lacked large parts of the sequence, one PLCP (ALP) carried a C-terminal tandem fusion, and one lacked the signal peptide required for targeting to the endomembrane system. Importantly, none of these mis-annotated proteins could have been active proteases, confirming that their annotation is incorrect. By contrast, the NbDE annotation would predict functional, active proteases that could react with activity-based probes and hence explain the purification of these proteins in activity profiling experiments.

### Two examples of improved subtilase annotations

To further illustrate gene annotations in the different datasets and show the relevance of adding missing genes from different genome assemblies with the NbE dataset, we analysed the gene-models of two subtilases which are missing in the Niben1.0.1 draft genome assembly. Both subtilases are encoded by a single-exon gene-models in the NbDE dataset (**Figure 4**). The sequence of NbE05066806.1 is present on three non-overlapping contigs in the Niben1.0.1 dataset, one of which contains three single nucleotide polymorphisms (SNPs, **Figure 4A**). Two of these SNPs are also present in the Niben0.4.4 dataset, but this annotation contains an additional 132bp insertion which is annotated as coding. The coding sequence of this subtilase is incomplete in the Nbv0.5 dataset and we identified 13 sequences corresponding to partial or chimeric variants of this subtilase in the Nb5.1 primary + alternate predicted proteome.

**Figure 4.**
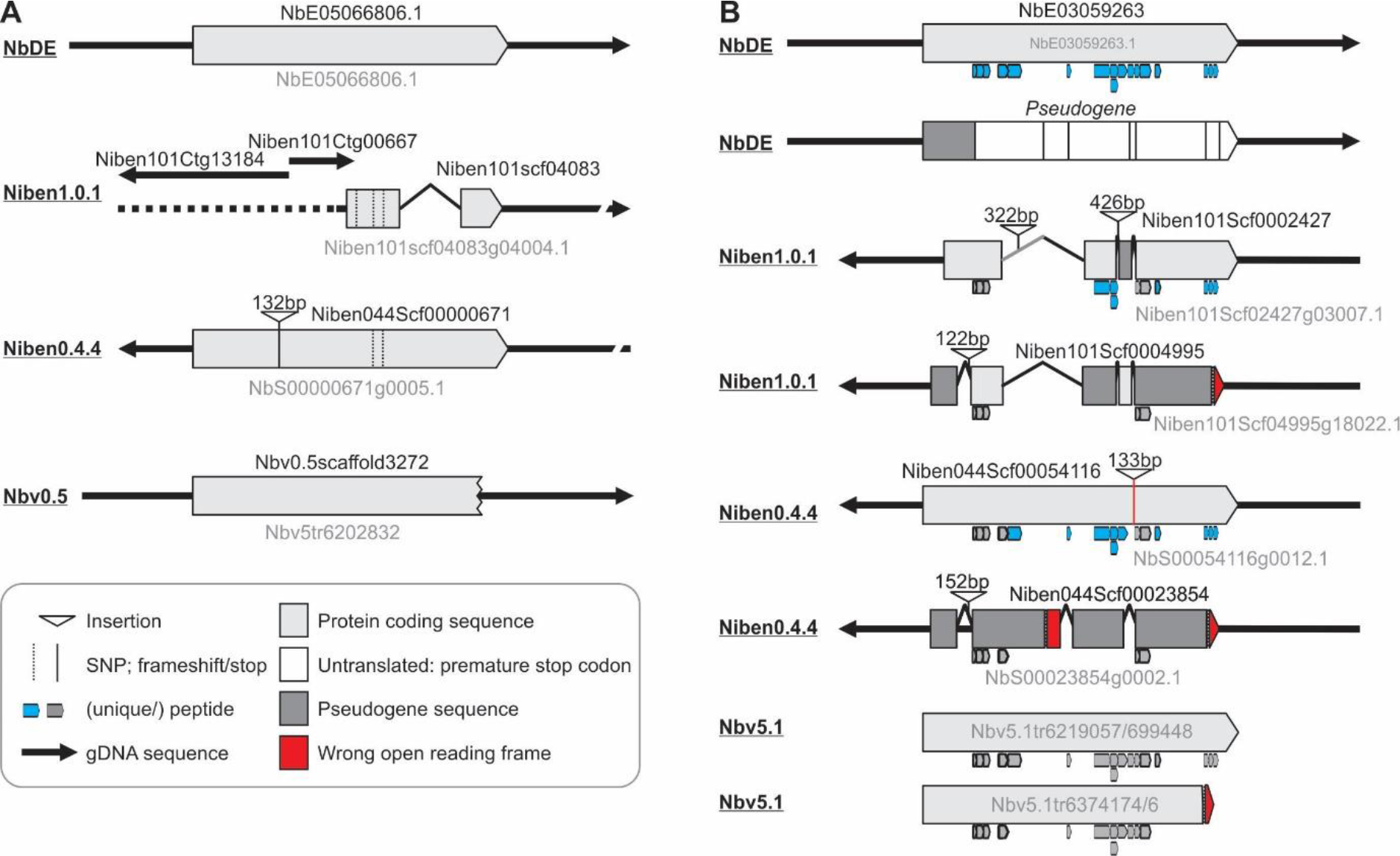
Examples of subtilase mis-annotations in the different genome assemblies. **(A)** Gene-models corresponding to subtilase NbE05066806 and the corresponding annotations in the various datasets. This subtilase gene is fragmented in Niben1.0.1; truncated in Nbv0.5; and carries two SNPs and an extra sequence in Niben0.4.4. **(B)** Gene-models corresponding to subtilase NbE03059263 and the corresponding annotations in the various datasets. This subtilase has a pseudogenised homeolog (dark grey) that was not retained in the NbDE dataset as it encodes a protein with <60% coverage because it contains premature stop codons. The pseudogene caused mis-assembly in the Niben1.0.1 dataset, resulting in a hybrid sequence. Mis-annotated exon-intron boundaries also effected gene models in Niben1.0.1, Niben0.4.4 and Nbv5.1. Peptides matched to the different gene models are indicated below the gene models.

A more complicated situation exists for subtilase NbE03059263, which we detected in the apoplast by proteomics. This subtilase has a close paralog which has been pseudogenised which was not retained in the NbDE annotation because it contains a premature stop codon and the coverage of the encoded protein is therefore below 60% (**Figure 4B**). Consequently, only NbE03059263 is detected with 17 unique peptides. In Niben1.0.1, however, sequences of these two homologous subtilases are misassembled, resulting in hybrids. In addition, several exon-intron boundaries have been misannotated, so the predicted proteins lack several crucial sequences. Consequently, only 12 of the peptides match the annotated subtilases in the Niben1.0.1 dataset, of which seven are unique. In Niben0.4.4, one subtilase gene carries a 133bp insertion that is annotated as an intron, but this insertion causes an amino acid substitution that prevents a match with one of the peptides. The paralog in Niben0.4.4 is mis-annotated as protein-encoding and consequently only 10 of the 16 peptides are unique. Finally, this subtilase is represented by four transcripts in the Nbv5.1 dataset, two of which encode a C-terminally truncated protein that lacks a match with three peptides, and none of the peptides are unique. Transcript sequences from the pseudogene are absent from this Nbv5.1 dataset. These two examples illustrate the different issues that occur with the gene annotations in the different datasets.

### Pseudogenization of subtilases is consistent with a contracting functional genome

To study pseudogenisation of homeologs further, we investigated the subtilase-encoding gene family. Several subtilisases are implicated in immunity, most notably the tomato P69 clade of subtilases (Taylor and Qiu, 2017). Our NbDE database contains 64 complete subtilase genes, and one partial gene. 16 of the subtilase genes are likely duplicated, and an additional 18 are putative pseudo-genes with >60% coverage to the NCBI *Nicotiana* RefSeq proteins. Three subtilases are missing in the NbD dataset, highlighting the utility of combining the NbD and NbE datasets to increase coverage of the proteome. Remarkably, no SBT3 clade family members were identified in *N. benthamiana* (**Table 1**, Supplemental **Figures S2-S3**). Three *N. benthamiana* subtilases may possess phytaspase activity based on the presence of an aspartic acid residue at the pro-domain junction as well as a histidine residue in the S1 pocket which is thought to bind to P1 aspartic acid (Supplemental **Figures S2-S3**, Reichardt et al., 2018).

**Table 1:**
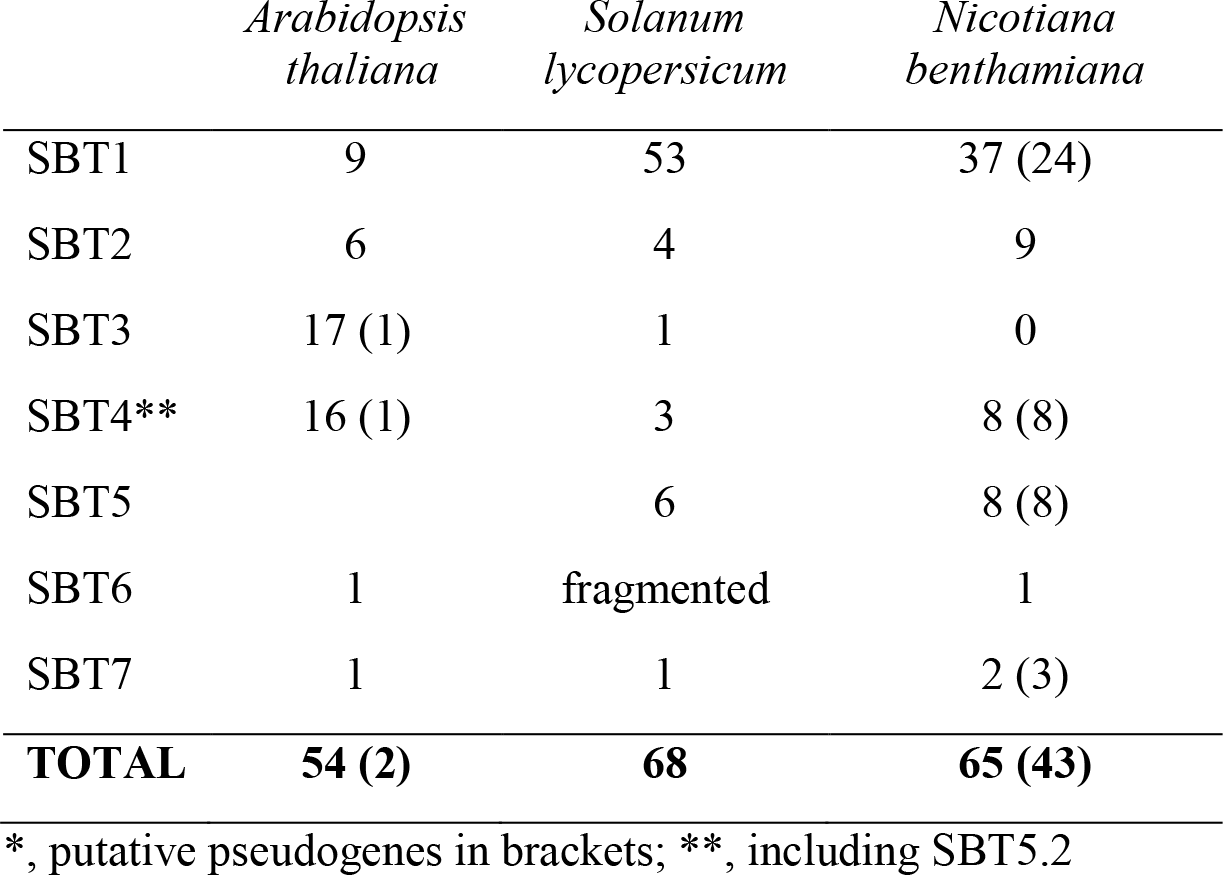
Subtilase gene number according to family*

In comparison, the Niben1.0.1 genome annotation predicts 103 different subtilase gene products. However, 38 of these genes are pseudo-genes and 16 subtilase genes are absent from Niben1.0.1 (Supplemental **Table S1** for a comprehensive comparison). Importantly, none of the remaining 49 subtilase-encoding genes in the Niben1.0.1 dataset are correctly annotated to encode a functional product. Furthermore, the predicted proteome from the Nbv5.1 primary + alternate transcriptome contains more than 400 subtilase gene products, largely due to a large number of chimeric sequences.

By searching the Niben1.0.1 and Niben0.4.4 genome assemblies using BLASTN, we identified 43 putative subtilase pseudo-genes, which had internal stop-codons and/or frame-shift mutations and are therefore likely non-coding. Phylogenetic analysis shows that these likely homeologs are often pseudogenized. This pattern of pseudogenization in the subtilase gene family is consistent with a contracting functional genome upon polyploidization, where for many functional protein-encoding gene there is a corresponding homeolog that is pseudogenized (**Figure 5**, **Table 1**, and Supplemental **Figure S2**). In conclusion, the new genome annotation represents a significant improvement over previous annotations and facilitates more accurate and meaningful phylogenetic analysis of gene families in *N. benthamiana*.

**Figure 5.**
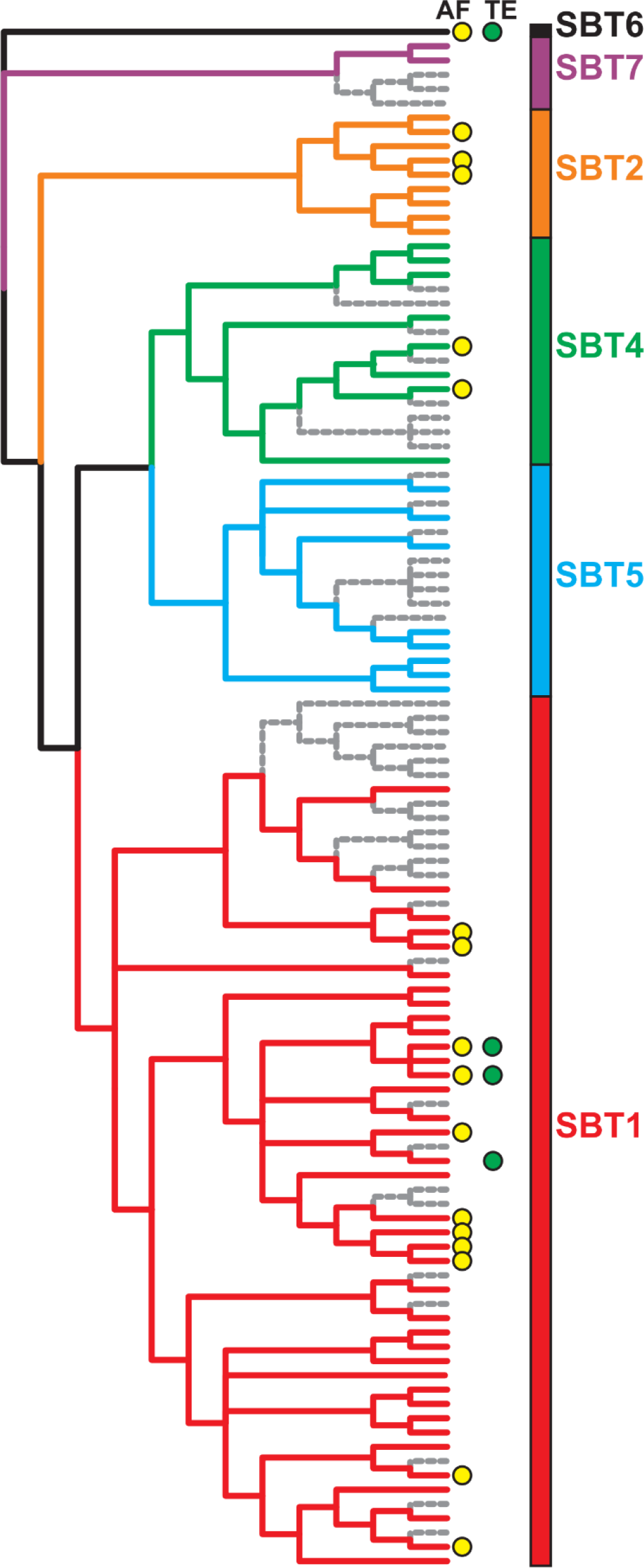
Birth and death of subtilase paralogs in *N. benthamiana*. The evolutionary history of the subtilase gene family was inferred by using the Maximum Likelihood method based on the Whelan and Goldman model. The bootstrap consensus tree inferred from 500 replicates is taken to represent the evolutionary history of the taxa analysed. Putative pseudogenes are indicated in grey. Subtilases identified in apoplastic fluid (AF) and/or total extract (TE) are indicated with yellow and green dots, respectively. Naming of subtilase clades is according to (Taylor and Qiu, 2017). Supplemental **Figure S2** includes the individual names.

### Hydrolases are enriched in the leaf secretome of *N. benthamiana*

Finally, we used the NbDE dataset to analyse the extracellular protein repertoire (secretome) of the *N. benthamiana* apoplast. The plant apoplast is the primary interface in plant-pathogen interactions (Misas-Villamil and Van der Hoorn, 2008; Doehlemann and Hemetsberger, 2013) and apoplastic proteins include many enzymes that may act in plant-pathogen interactions. To identify apoplastic proteins, we performed MS analysis of apoplastic fluids (AFs) and total extracts (TEs) isolated from the same leaves in four biological replicates. Annotation using the NbDE dataset show that these AF and TE proteomes are clearly distinct (**Figure 6A**). We assigned 510 proteins to the apoplast because they were only detected in AF samples or highly enriched in the AF samples over TE samples (log_2_ ≥1.5, p ≤0.01, BH-adjusted moderated t-test, **Figure 6B**). Similarly, we assigned 1042 intracellular proteins because they were only detected in the TE samples or enriched in TE samples over AF samples (log_2_ ≤−1.5, p≤0.01, **Figure 6B**). The remaining 833 proteins were considered both apoplastic and intracellular. As expected, the apoplastic proteome is significantly enriched for proteins containing a SignalP-predicted signal peptide, while the intracellular proteins and proteins present both in the apoplast and intracellular are significantly enriched for proteins lacking a signal peptide (BH-adjusted hypergeometric test, p<0.001).

**Figure 6.**
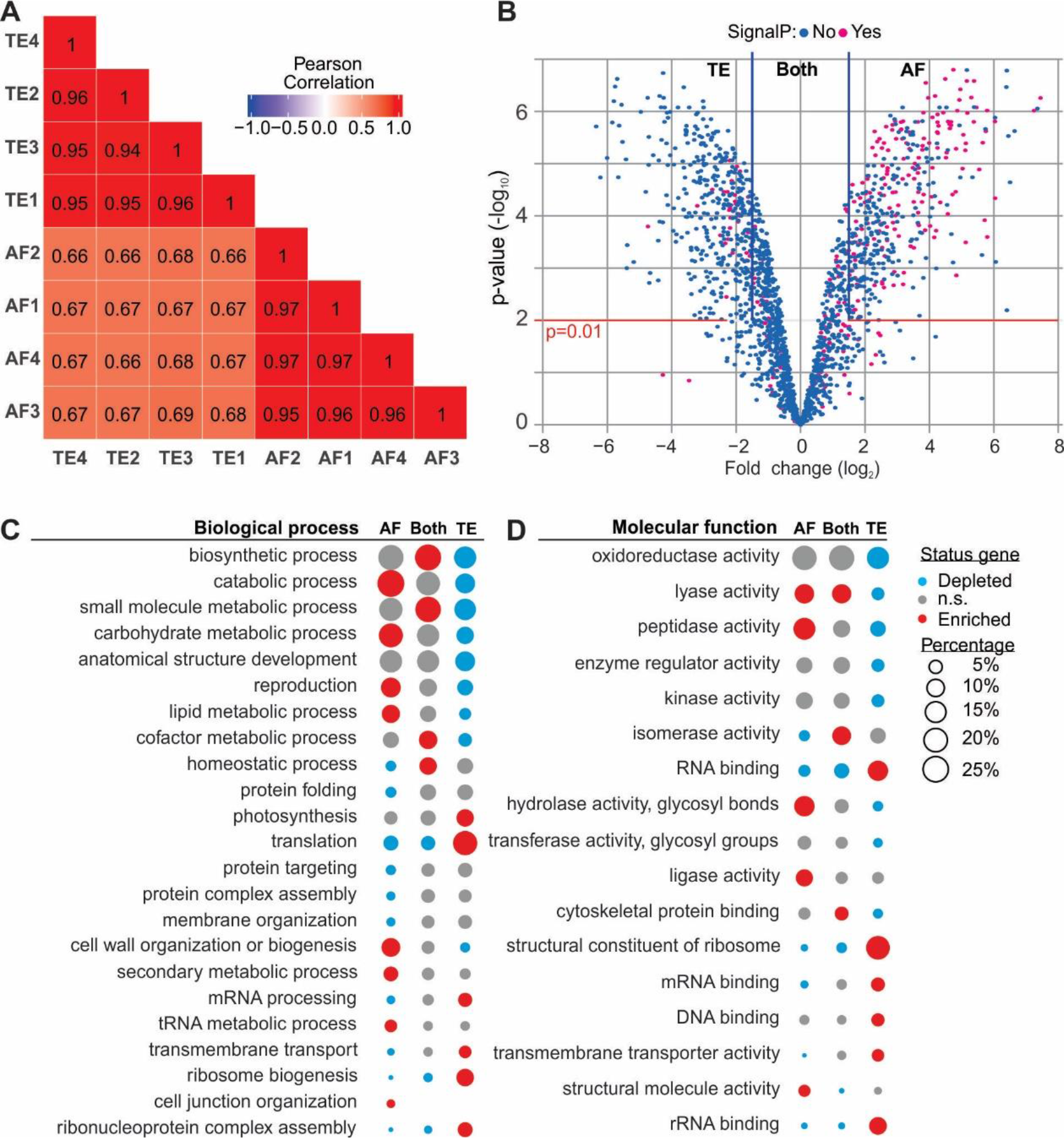
Annotation of the *N. benthamiana* apoplastic proteome. **(A)** Correlation matrix heat map of the log_2_ transformed LFQ intensity of protein groups in the four biological replicates of apoplastic fluid (AF) and total extract (TE) samples. Biological replicates are clustered on similarity. **(B)** A volcano plot is shown plotting log_2_ fold difference of AF/TE over −log_10_ BH-adjusted moderated p-values. Proteins with log_2_ ≥1.5 and p≤0.01 were considered apoplastic, as well as those only found in AF. Conversely, proteins with a log_2_ ≤1.5 and p≤0.01 were considered intracellular, as well as those found only in TE. **(C)** Percentage of proteins in each fraction annotated with biological process-associated GO-SLIM terms. **(D)** Percentage of proteins in each fraction annotated with molecular function-associated GO-SLIM terms. **(C-D)** GO-SLIM annotations are shown when significantly enriched or depleted (BH-adjusted hypergeometric test, p<0.05) in at least one of the fractions (AF, TE, or both). Each bubble indicates the percentage of genes containing that specific GO-SLIM annotation in that compartment. Colours indicate whether the GO-SLIM annotations are enriched or depleted in that fraction (p<0.05, n.s., non-significant).

The apoplastic proteome is enriched for proteins acting in catabolic processes and carbohydrate and lipid metabolic processes (**Figure 6C**), which is reflected in the enrichment of peptidases, glycosidases and other hydrolases (**Figure 6D**, Supplemental **Table S2** for a full list). Proteins considered predominantly intracellular are enriched for GO-SLIM terms associated with translation, photosynthesis and transport as biological processes (**Figure 6C**), and a similar pattern is seen for GO-SLIM terms describing molecular functions (**Figure 6D**). Proteins present both in TE and in AF are enriched for GO-SLIM terms associated with biosynthetic processes and homeostasis (**Figure 6C**). These processes are usually performed by proteins acting at multiple subcellular localizations.

To specify which peptidases are enriched in the apoplast, we assigned the PFAM-annotations to MEROPS peptidase identifiers (Rawlings *et al.*, 2018). Three of the 15 different families of peptidases detected in the apoplast have significantly more members enriched in the AF as compared to TE: the subtilase family (S08; 13 members, p<0.001), serine carboxypeptidase-like family (S10; 8 members, p<0.01), and aspartic peptidase family (A01; 17 members, p<0.001). By contrast, the proteasome is enriched in the intracellular fraction (T01; 26 members, p<0.001) (BH-adjusted hypergeometric test, Supplemental **Table S3** for a full list).

### Conclusions and prospects

Homology-guided re-annotation of the *Nicotiana benthamiana* genome resulted in improved gene models for the alloploid model plant *Nicotiana benthamiana*. This approach identifies many genes missing from previous annotations, and improves the annotation of exon-intron boundaries and overcomes problems associated with mis-assignment of short reads from pseudogenized homeologs. By removing sequences with <60% coverage to well-annotated *Nicotiana* proteins, we removed partial sequences and products of pseudogene products caused by premature stop codons and frameshift mutations. Besides the core NbD dataset containing 53,411 coding sequences, we provide a supplemental NbE dataset with 21,391 coding sequences, including homeologs. Both the core NbD dataset and the combined NbDE datasets have longer protein sequences with increased BUSCO scores and higher coverage to the tomato proteome. These datasets also have improved frequency of PFAM annotations while maintaining a relatively low number of entries. Both datasets outperform the preceding datasets in the annotation of spectra during proteomics experiments. These datasets provide the research community with improved capacity to annotate spectra during proteomics experiments. These datasets will also provide a valuable basis for further genome annotation and reverse genetic approaches in *Nicotiana benthamiana*.

### Material & Methods

#### Sequence retrieval

**Table 2** summarizes the genomes and genome annotations used in this study.

**Table 2:** Genomes and gene annotations used*

| Species | Genome build | Annotation | Reference |
| --- | --- | --- | --- |
| *Arabidopsis thaliana* ecotype Colombia | GCF_000001735.4 | RefSeq | Arabidopsis Genome Initiative, 2000 |
| *Beta vulgaris* subsp. *vulgaris* | GCF_000511025.2 | NCBI *Beta vulgaris* subsp. *vulgaris* Annotation Release 101 | Dohm et al., 2014 |
| *Capsicum annuum* cv. CM334 | GCA_000512255.2 | Pepper.v.1.55.proteins.annotated | Qin et al., 2014 |
| *Capsicum annuum* cv. Zunla-1 | GCF_000710875.1 | NCBI *Capsicum annuum* Annotation Release 100 | Kim et al., 2014 |
| *Capsicum annuum* var. *glabriusculum* | GCA_000950795.1 | CaChiltepin.pep | Qin et al., 2014 |
| *Daucus carota* subsp. *sativus* cv. DH1 | GCF_001625215.1 | NCBI *Daucus carota* subsp. *sativus* Annotation Release 100 | Iorizzo et al., 2016 |
| *Nicotiana attenuata* strain UT | GCF_001879085.1 | NCBI *Nicotiana attenuata* Annotation Release 100 | Xu et al., 2017 |
| *Nicotiana benthamiana* | Niben1.0.1 | Niben101_annotation.proteins.wdesc | Bombarely et al., 2012 |
| *Nicotiana benthamiana* | Niben0.4.4 | Niben.genome.v0.4.4.proteins.wdesc | Bombarely et al., 2012 |
| *Nicotiana benthamiana* | Nbv0.3 | - | Naim et al., 2012 |
| *Nicotiana benthamiana* | Nbv0.5 | - | Naim et al., 2012 |
| *Nicotiana benthamiana* | Nbv5.1 transcriptome | Nbv5.1_transcriptome_primary_alternate_correct | Nakasugi et al., 2014 |
| *Nicotiana obtusifolia* cv. 1x inbred | GCA_002018475.1 | NIOBT_r1.0 | Xu et al., 2017 |
| *Nicotiana sylvestris* | GCF_000393655.1 | NCBI *Nicotiana sylvestris* Annotation Release 100 | Sierro et al., 2013 |
| *Nicotiana tabacum* cv. TN90 | GCF_000715135.1 | NCBI *Nicotiana tabacum* Annotation Release 100 | Sierro et al., 2014 |
| *Nicotiana tomentosiformis* | GCF_000390325.1 | NCBI *Nicotiana tomentosiformis* Annotation Release 101 | Sierro et al., 2013 |
| *Petunia axillaris* N | Petunia_axillaris_v1.6.2 | Petunia_axillaris_v1.6.2_proteins | Bombarely et al., 2016 |
| *Petunia inflata* S6 | Petunia_inflata_v1.0.1 | Petunia_inflata_v1.0.1_proteins | Bombarely et al., 2016 |
| *Solanum lycopersicum* cv. Heinz 1706 | GCF_000188115.4 | NCBI *Solanum lycopersicum* Annotation Release 103 | Tomato Genome Consortium, 2012 |
| *Solanum melongena* L. cv. Nakate-Shinkuro | GCA_000787875.1 | SME_r2.5.1_pep | Hirakawa et al., 2014 |
| *Solanum pennellii* | GCF_001406875.1 | NCBI *Solanum pennellii* Annotation Release 101 | Bolger et al., 2014 |
| *Solanum tuberosum* cv. DM 1-3 516 R44 | GCF_000226075.1 | NCBI *Solanum tuberosum* Annotation Release 101 | Potato Genome Sequencing Consortium et al., 2011 |

\*, Where available the NCBI assembly accession and annotation was taken

#### Annotation

In order to extract gene-models from the published *N. benthamiana* draft genomes we combined NCBI *Nicotiana* RefSeq protein sequences in one database, removed all partial proteins and those containing undetermined residues, added 110 genes which we had previously been manually curated and 565 full-length *N. benthamiana* proteins from GenBank leading to a dataset containing 226,543 protein sequences. We used CD-HIT (v4.6.8) (Fu *et al.*, 2012) to cluster these sequences at a 95% identity threshold and reduce the redundancy in our database while removing partials (Nicotiana_db95; 85,453 sequences). This dataset was used to annotate the gene-models in the different *N. benthamiana* genome builds using Scipio version 1.4.1 (Keller *et al.*, 2008) which was run to allow for duplicated genes. After running Scipio we used Augustus (v3.3) (Stanke *et al.*, 2006) to extract complete and partial gene models. Transdecoder (v5.5.0) (Haas *et al.*, 2013) was used to retrieve the single-best ORF containing homology to the Nicotiana_db95 database as determined by BLASTP for gene models containing internal stop codons. The predicted proteins were aligned to the NCBI *Nicotiana* RefSeq protein sequences using BLASTP (NCBI-BLAST v2.8.1+), and only those with a coverage ≥ 60% in the Niben1.0.1 genome assembly, or ≥ 90% in the Niben0.4.4, Nbv0.3, and Nbv0.5 genome assemblies were maintained. A custom R script was used to convert extracted Niben1.0.1 gene models in a full genome annotation, which were manually inspected resulting in **NbD**. CD-HIT-2D was used at 100% identity to identify gene models missing in NbD, but present in the extracted gene models from the other genome assemblies (the **NbE** set). The addition of these sequences to NbD resulted in the **NbDE** annotation. This dataset was annotated using SignalP (v4.1) (Petersen *et al.*, 2011), PFAM (v32) (El-Gebali *et al.*, 2019) and annotated the with GO terms and UniProt identifiers using Sma3s v2 (Casimiro-Soriguer *et al.*, 2017).

We ran BUSCO (v3.0.2; dependencies: NCBI-BLAST v2.8.1+ HMMER v3.1; Augustus v3.3) (Simão *et al.*, 2015; Waterhouse *et al.*, 2018) on the different *N. benthamiana* predicted proteomes using the plants set (Embryophyta_odb9), to validate their completeness. Additionally, we used BLASTP to compare these proteins with the NCBI *Solanum lycopersicum* RefSeq protein sequences.

#### Sample Preparation For Proteomics And Definition Of Biological Replicates

Four-week old *N. benthamiana* plants were used. The AF was extracted by vacuum infiltrating *N. benthamiana* leaves with ice-cold MilliQ. Leaves were dried to remove excess liquid, and apoplastic fluid was extracted by centrifugation of the leaves in a 20 ml syringe barrel (without needle or plunger) in a 50 ml falcon tube at 2000 × g, 4°C for 25 min. Samples were snap-frozen in liquid nitrogen and stored at −80°C prior to use. TE was collected by removing the central vein and snap-freezing the leaves in liquid nitrogen followed by grinding in a pestle and mortar and addition of three volumes of phosphate-buffered saline (PBS) (w/v). One biological replicate was defined as a sample, AF or TE, consisting of one leaf from three independent plants (3 leaves total). Four independent biological replicates were taken for AF and TE.

#### Protein digestion and sample clean-up

AF and TE sample corresponding to 15 μg of protein was taken for each sample (based on Bradford assay). Dithiothreitol (DTT) was added to a concentration of 40 mM, and the volume adjusted to 250 μl with MS-grade water (Sigma). Proteins were precipitated by the addition of 4 volumes of ice-cold acetone, followed by a 1 hr incubation at −20°C and subsequent centrifugation at 18,000 × g, 4°C for 20 min. The pellet was dried at room temperature (RT) for 5 min and resuspended in 25 µL 8 M urea, followed by a second chloroform/methanol precipitation. The pellet was dried at RT for 5 min and resuspended in 25 µL 8 M urea. Protein reduction and alkylation was achieved by sequential incubation with DTT (final 5 mM, 30 min, RT) and iodoacetamide (IAM; final 20 mM, 30 min, RT, dark). Non-reacted IAM was quenched by raising the DTT concentration to 25 mM. Protein digestion was started by addition of 1000 ng LysC (Wako Chemicals GmbH) and incubation for 3 hr at 37°C while gently shaking (800 rpm). The samples were then diluted with ammoniumbicarbonate (final concentration 80 mM) to a final urea concentration of 1 M. 1000 ng Sequencing grade Trypsin (Promega) was added and the samples were incubated overnight at 37°C while gently shaking (800 rpm). Protein digestion was stopped by addition of formic acid (FA, final 5% v/v). Tryptic digests were desalted on home-made C18 StageTips (Rappsilber *et al.*, 2007) by passing the solution over 2 disc StageTips in 150 µL aliquots by centrifugation (600-1200 × g). Bound peptides were washed with 0.1 % FA and subsequently eluted with 80% Acetonitrile (ACN). Using a vacuum concentrator (Eppendorf) samples were dried, and the peptides were resuspended in 20 µL 0.1% FA solution.

#### LC-MS/MS

The samples were analysed as in (Grosse-Holz *et al.*, 2018). Briefly, samples were run on an Orbitrap Elite instrument (Thermo) (Michalski *et al.*, 2011) coupled to an EASY-nLC 1000 liquid chromatography (LC) system (Thermo) operated in the one-column mode. Peptides were directly loaded on a fused silica capillary (75 µm × 30 cm) with an integrated PicoFrit emitter (New Objective) analytical column packed in-house with Reprosil-Pur 120 C18-AQ 1.9 µm resin (Dr. Maisch), taking care to not exceed the set pressure limit of 980 bar (usually around 0.5-0.8 µl/min). The analytical column was encased by a column oven (Sonation; 45°C during data acquisition) and attached to a nanospray flex ion source (Thermo). Peptides were separated on the analytical column by running a 140 min gradient of solvent A (0.1% FA in water;; Ultra-Performance Liquid Chromatography (UPLC) grade) and solvent B (0.1% FA in ACN; UPLC grade) at a flow rate of 300 nl/min (gradient: start with 7% B; gradient 7% to 35% B for 120 min; gradient 35% to 100% B for 10 min and 100% B for 10 min) at a flow rate of 300 nl/min.). The mass spectrometer was operated using Xcalibur software (version 2.2 SP1.48) in positive ion mode. Precursor ion scanning was performed in the Orbitrap analyzer (FTMS; Fourier Transform Mass Spectrometry) in the scan range of m/z 300-1800 and at a resolution of 60000 with the internal lock mass option turned on (lock mass was 445.120025 m/z, polysiloxane) (Olsen *et al.*, 2005). Product ion spectra were recorded in a data-dependent manner in the ion trap (ITMS) in a variable scan range and at a rapid scan rate. The ionization potential was set to 1.8 kV. Peptides were analysed by a repeating cycle of a full precursor ion scan (1.0 × 106 ions or 50ms) followed by 15 product ion scans (1.0 × 10^4^ ions or 50ms). Peptides exceeding a threshold of 500 counts were selected for tandem mass (MS2) spectrum generation. Collision induced dissociation (CID) energy was set to 35% for the generation of MS2 spectra. Dynamic ion exclusion was set to 60 seconds with a maximum list of excluded ions consisting of 500 members and a repeat count of one. Ion injection time prediction, preview mode for the Fourier transform mass spectrometer (FTMS, the orbitrap), monoisotopic precursor selection and charge state screening were enabled. Only charge states higher than 1 were considered for fragmentation.

#### Peptide and Protein Identification

Peptide spectra were searched in MaxQuant (version 1.5.3.30) using the Andromeda search engine (Cox *et al.*, 2011) with default settings and label-free quantification and match-between-runs activated (Cox and Mann, 2008; Cox *et al.*, 2014) against the databases specified in the text including a known contaminants database. Included modifications were carbamidomethylation (static) and oxidation and N-terminal acetylation (dynamic). Precursor mass tolerance was set to ±20 ppm (first search) and ±4.5 ppm (main search), while the MS/MS match tolerance was set to ±0.5 Da. The peptide spectrum match FDR and the protein FDR were set to 0.01 (based on a target-decoy approach) and the minimum peptide length was set to 7 amino acids. Protein quantification was performed in MaxQuant (Tyanova *et al.*, 2016), based on unique and razor peptides including all modifications.

#### Proteomics processing in R

Identified protein groups were filtered for reverse and contaminants proteins and those only identified by matching, and only those protein groups identified in 3 out of 4 biological replicates of either AF or TE were selected. The LFQ values were log_2_ transformed, and missing values were imputed using a minimal distribution as implemented in imputeLCMD (v2.0) (Lazar, 2015). A moderated t-test was used as implemented in Limma (v3.34.3) (Ritchie *et al.*, 2015; Phipson *et al.*, 2016) and adjusted using Benjamini–Hochberg (BH) adjustment to identify protein groups significantly differing between AF and TE. Bonafide apoplastic protein groups were those only detected in AF and those significantly (p≤0.01) log_2_ fold change ≥1.5 in AF samples. Protein groups only detected in TE and those significantly (p≤0.01) log_2_ fold change ≤−1.5 depleted in AF samples were considered intracellular. The remainder was considered both apoplastic and intra-cellular. Majority proteins were annotated with SignalP, PFAM, MEROPS (v12) (Rawlings *et al.*, 2018), GO, and UniProt keywords identifiers. A BH-adjusted Hypergeometric test was used to identify those terms that were either depleted or enriched (p≤0.05) in the bonafide AF protein groups as compared to bonafide AF depleted proteins or protein groups present both in the AF and TE.

#### Phylogenetic analysis

Predicted proteomes were annotated with PFAM identifiers, and all sequences containing a Peptidase S8 (PF00082) domain were extracted from the different databases. Additionally, we manually curated the subtilase gene-family in the Niben1.0.1 draft genome, identifying putative pseudo-genes which were annotated as protein-encoding genes, as well as missing genes and incorrect gene models or genes in which the reference sequence was absent in Niben1.0.1. Tomato subtilases were retrieved from Solgenomics, and other previously characterized subtilases (Taylor and Qiu, 2017) were retrieved from NCBI. Clustal Omega (Sievers *et al.*, 2011; Li *et al.*, 2015) was used to align these sequences. The putative pseudo-gene sequences were substituted with the best blast hit in NCBI to visualize pseudogenization in the alignment and phylogenetic tree. Determining the best model for maximum likelihood phylogenetic analysis and the phylogenetic analysis was performed in MEGA X (Kumar *et al.*, 2018). The evolutionary history was inferred by using the Maximum Likelihood method based on the Whelan and Goldman model. A discrete Gamma distribution was used to model evolutionary rate differences among sites, and the rate variation model allowed for some sites to be evolutionarily invariable. All positions with less than 80% site coverage were eliminated. Niben101Scf00595_742942-795541 was used to root the phylogenetic trees.

## Acknowledgements

We would like to thank Philippe Varennes-Jutras and Daniela Sueldo for critically reading the manuscript and providing important suggestions for improving the manuscript.

## Author contributions

Conceptualization: JK, RvdH; Formal analysis: JK; Funding acquisition: RvdH; Wetlab experiments: FHG; Proteomics: FK, MK; Programming: JK, FH; Writing: JK, RvdH.

---

**Figure S1:** Comparison of Solanaceae proteomes.

**Figure S2:** Phylogenetic analysis of the subtilase gene-family with names.

**Figure S3:** Phylogenetic analysis of the subtilase gene-family of tomato and Arabidopsis and including other previously characterized subtilases.

**Table S1:** Gene-model comparison

**Table S2:** GO-SLIM term enrichment complete at p≤0.05

**Table S3:** MEROPS family term enrichment complete

**Supplemental dataset 1**: New Niben1.0.1 gff3 annotation

**Supplemental dataset 2**: FASTA file of NbE genomic sequence ±1kb

**Supplemental dataset 3:** gff3 annotation of NbE gene-models

**Supplemental dataset 4**: NbD proteome

**Supplemental dataset 5**: NbD transcriptome

**Supplemental dataset 6**: NbDE proteome

**Supplemental dataset 7**: NbDE transcriptome

**Supplemental dataset 8**: Sma3s v2 annotation of NbDE

**Supplemental dataset 9**: PFAM32 annotation of NbDE

**Supplemental dataset 10**: SignalP4.1 annotation of NbDE

