## Supplemental Figures S1, S2, S3 for "Homology-guided re-annotation improves the gene models of the alloploid *Nicotiana benthamiana*"

**Supplemental figures** Kourelis et al: *Re-annotation of N. benthamiana gene models.*

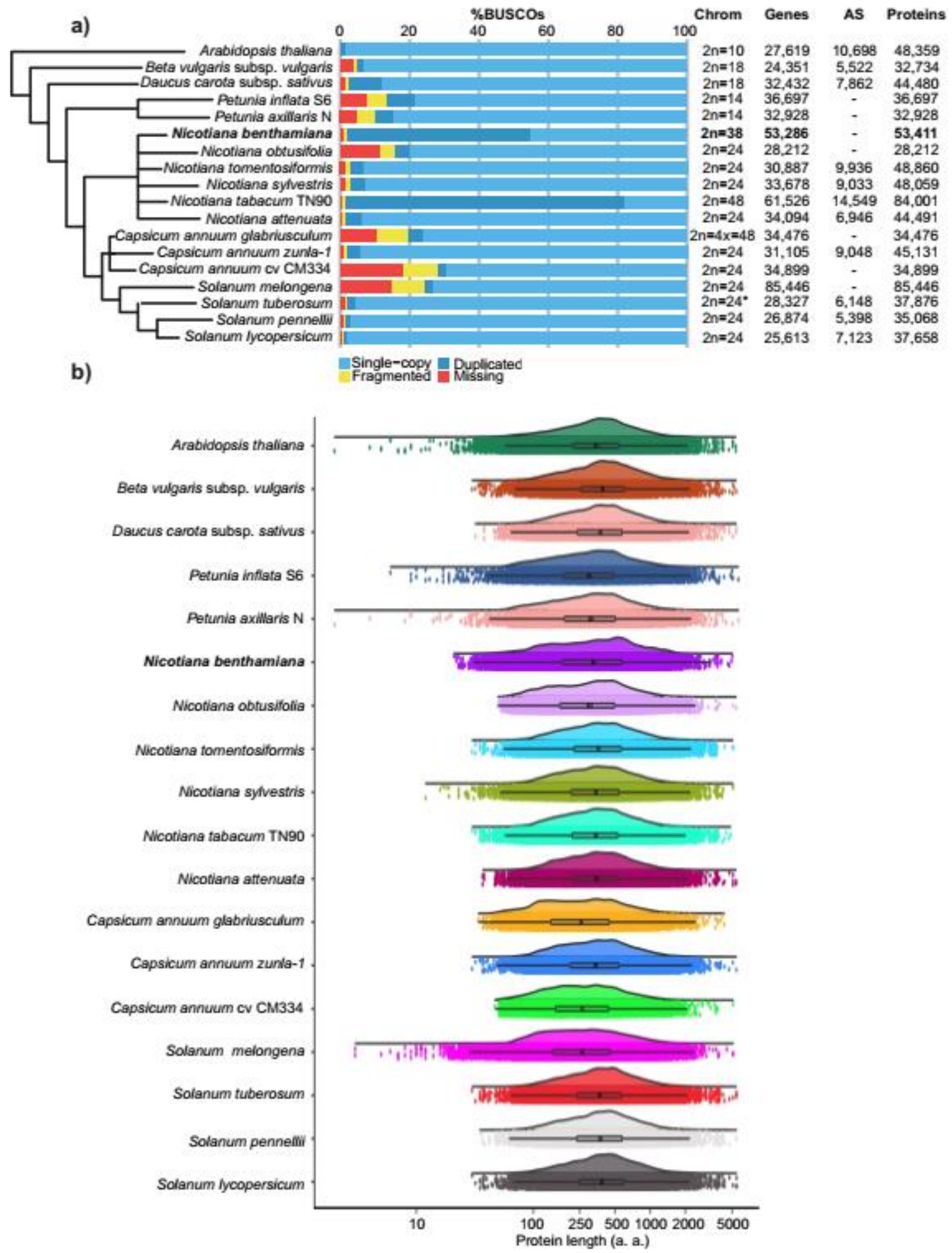


**Figure S1:** Comparison of Solanaceae proteomes. **a)** Completeness of the predicted proteomes from sequenced Solanaceae genomes was estimated using BUSCO v3 with the embryophyta database. *Certain *Solanum tuberosum* species are polyploid. **b)** Violin and boxplot g of $\log_{10}$ protein length distribution of each predicted proteome. Jittered dots show the raw underlying data. *Nicotiana benthamiana* is represented by the NbD dataset.

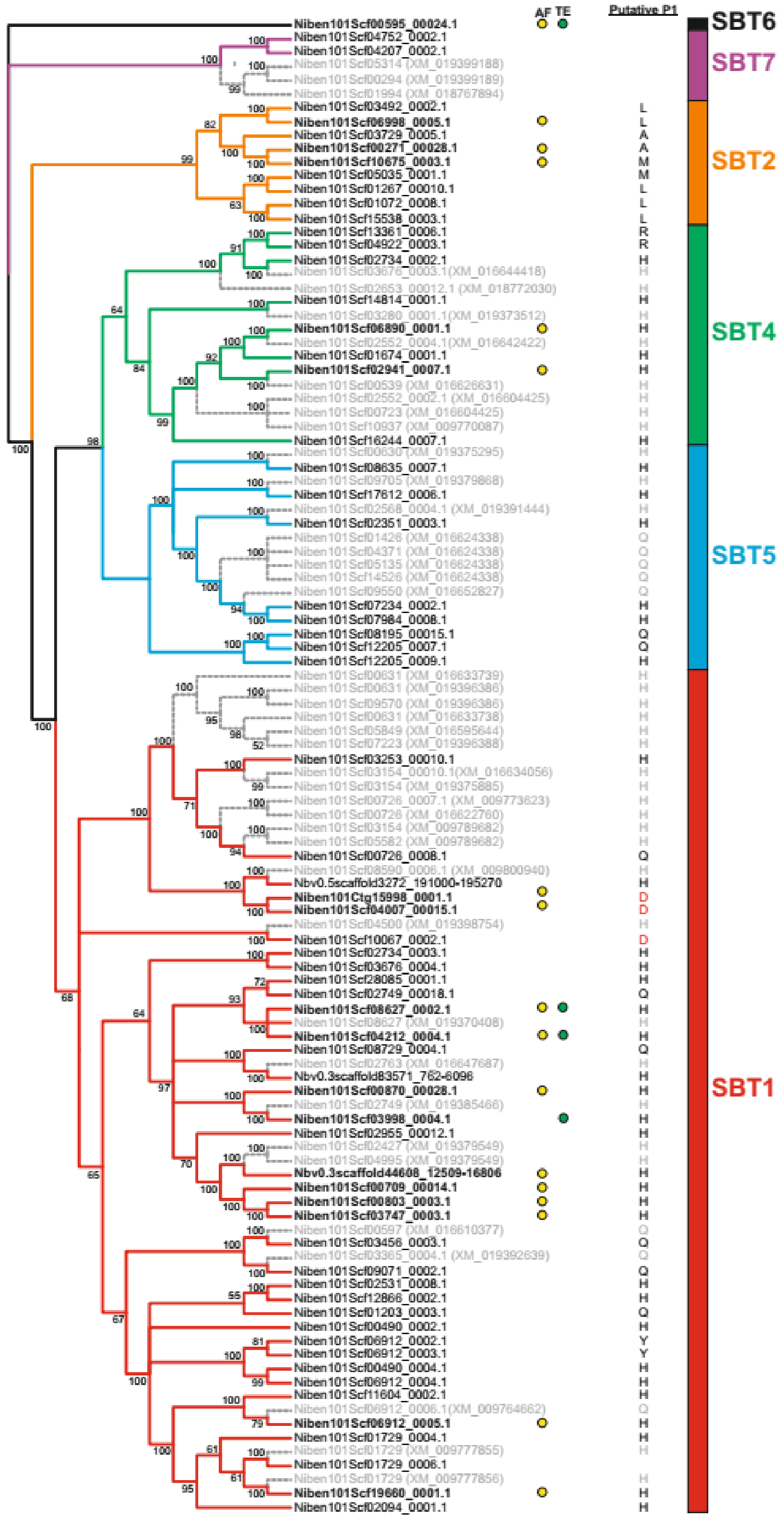


**Figure S2:** Phylogenetic analysis of the subtilisin gene-family with full names. The evolutionary history of the subtilase gene family was inferred by using the Maximum Likelihood method based on the Whelan and Goldman model. The bootstrap consensus tree inferred from 500 replicates is taken to represent the evolutionary history of the taxa analysed. Putative pseudogenes are indicated in grey. Subtilases identified in apoplastic fluid (AF) and/or total extract (TE) are indicated with yellow and green dots, respectively. Putative P1 based on residue at the prodomain junction is indicated, putative phytaspases are indicated in red. Naming of subtilase clades according to (Taylor and Qiu, 2017).

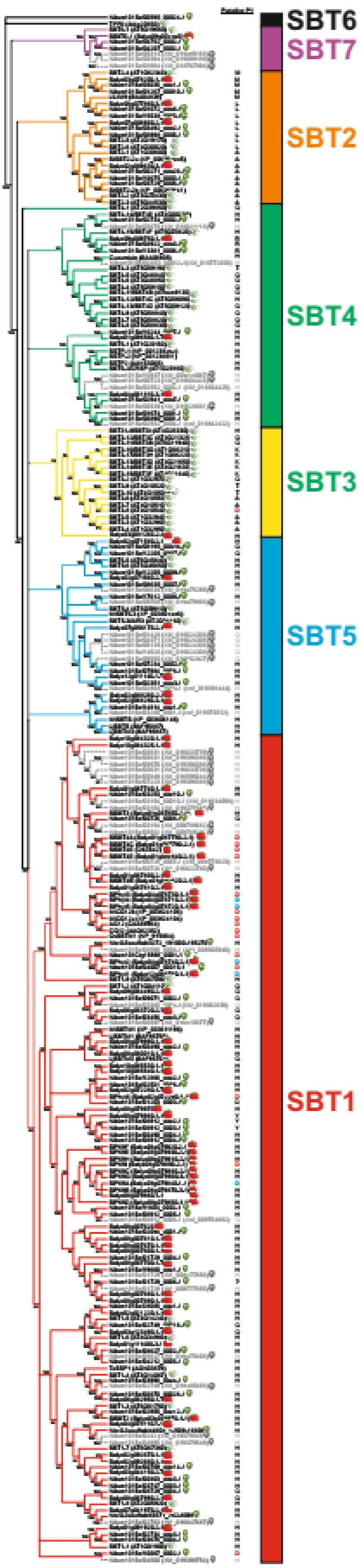


**Figure S3:** Phylogenetic analysis of the subtilisin gene-family of tomato and Arabidopsis and including other previously characterized subtilisins. The evolutionary history of the subtilase gene family was inferred by using the Maximum Likelihood method based on the Whelan and Goldman model. The bootstrap consensus tree inferred from 250 replicates is taken to represent the evolutionary history of the taxa analysed. Putative pseudogenes are indicated in grey. Putative P1 based on residue at the prodomain junction is indicated. Putative phytaspases are indicated in red, confirmed phytaspases in blue. Naming of subtilase clades according to (Taylor and Qiu, 2017).

### Other supplemental datasets:

**Table S1:** Gene-model comparison of subtilases

**Table S2:** GO-SLIM term enrichment complete at $p \leq 0.05$

**Table S3:** MEROPS family term enrichment complete

**Supplemental dataset 1**: New Niben1.0.1 gff3 annotation

**Supplemental dataset 2**: FASTA file of NbE genomic sequence ±1kb

**Supplemental dataset 3:** gff3 annotation of NbE gene-models

**Supplemental dataset 4**: NbD proteome

**Supplemental dataset 5**: NbD transcriptome

**Supplemental dataset 6**: NbDE proteome

**Supplemental dataset 7**: NbDE transcriptome

**Supplemental dataset 8**: Sma3s v2 annotation of NbDE

**Supplemental dataset 9**: PFAM32 annotation of NbDE

**Supplemental dataset 10**: SignalP4.1 annotation of NbDE
